# Mitochondrial Genome of the lesser known - Madras Hedgehog (*Paraechinus nudiventris*): Genomic characterization and comparative analysis within Erinaceidae

**DOI:** 10.1101/2025.05.13.653780

**Authors:** Thapasya Vijayan, Harald Meimberg, Brawin Kumar

## Abstract

The Madras hedgehog (*Paraechinus nudiventris*) is an understudied species endemic to southern India, with limited genetic data available for understanding its evolutionary history. Here, we present the complete mitochondrial genome (17,232 bp), annotated and analyzed in comparison with other *Erinaceidae* species. The mitogenome retains the typical vertebrate structure, comprising 14 protein-coding genes, 22 tRNAs, 2 rRNAs, and a non-coding control region, with an A+T rich nucleotide composition. Phylogenetic analyses using Bayesian and maximum likelihood approaches confirmed the species’ placement within *Paraechinus*, with divergence from *P. micropus* estimated during the Pleistocene. Molecular evolutionary analyses revealed selective constraints on oxidative phosphorylation genes, with *atp8* showing the highest variability. Our divergence time estimates, incorporating fossil calibrations, refine the evolutionary timeline of *Erinaceidae*, providing insights into the diversification of hedgehogs in South Asia. By making this annotated mitochondrial genome publicly available, we provide a foundational resource for future studies on the genetic diversity, phylogeography, and conservation of *P. nudiventris* and its relatives. The study also demonstrates the utility of non-invasive sampling methods in generating high-quality genomic data for cryptic mammalian species.

## Introduction

The Madras Hedgehog (*Paraechinus nudiventris*) endemic to southern India, is known for its highly cryptic behavior, nocturnal habits, and ability to thrive in urbanized landscapes – an extraordinary feat for a species typically associated with remote habitats. Unlike many other hedgehog species that are confined to forested or rural areas, the Madras Hedgehog has adapted to living in the increasingly urbanized areas of Tamil Nadu (Kumar et al. 2019b), where it is often spotted in city outskirts, agricultural land, and even rocky urban areas. Despite its widespread presence in certain regions, its elusive nature and nocturnal habits make studying its population dynamics and ecological requirements a considerable challenge. The species’ ability to navigate anthropogenic landscapes places it in a unique ecological niche, yet it remains largely under-studied in terms of genetics and evolutionary biology. The Madras hedgehog is one of five hedgehog species in India, and although it is classified as ‘Least Concern’ by the IUCN (Molur et al. 2005), there is a pressing need for genetic studies to understand its evolutionary relationships and conservation requirements. Anthropogenic pressures, including habitat loss and urbanization, pose significant threats to its survival (Kumar et al. 2019a; Kumar et al. 2019b) making it imperative to assess its distribution and conservation status. Despite this status, habitat loss from fuel wood collection, agriculture, trade, pet and urbanization poses significant threats (Kumar et al. 2019b). Other major threats to their population include hunting, roadkill, and local trade (Kumar et al. 2019a; Kumar et al. 2019b). Limited baseline data, combined with the species’ cryptic behavior, have made it difficult to assess its true population distribution and status.

The Eulipotyphla, order, to which the hedgehog belongs, has a rich evolutionary history, with members of the Erinaceidae family dating back over 85.8 million years (Meredith et al. 2011). The order comprises insectivorous mammals from four families: Erinaceidae (hedgehogs and gymnures), Talpidae (moles), Soricidae (shrews), and Solenodontidae (solenodons). Currently, 540 species are recognized (Burgin et al. 2020), but many cryptic species are likely yet to be discovered, particularly in areas outside established protected regions (Kennerley et al. 2021; Parsons et al. 2022). The Erinaceidae family includes 27 living species categorized into two subfamilies: Erinaceinae, which comprises spiny hedgehogs, and Galericinae, which consists of silky-furred gymnures and moonrats (Frost et al. 1991). Within the Erinaceidae family, hedgehogs have evolved distinct traits such as their spiny protection and fossorial lifestyle. Hedgehogs are primarily native to Africa and Eurasia, with some species having wide distribution ranges (Corbet 1988; He et al. 2012). South Asia inhabits five hedgehog species, including *Hemiechinus auritus, Hemiechinus collaris, Paraechinus hypolmelas, Paraechinus micropus*, and *Paraechinus nudiventris* (Johnsingh and Po 2015). Despite the number of studies dedicated to hedgehog phylogeny, much of the focus has been on more widespread species, leaving *Paraechinus nudiventris* understudied in terms of molecular and genetic data. Recent work by (Zeng et al. 2024) has included *P*.*nudiventris* in a mitogenomic phylogenetic analysis, providing its first molecular data and revealing its evolutionary position within Paraechinus. However, their focus was on broader phylogenetic relationships across Erinaceidae, leaving significant gaps in the detailed genomic characterization and evolutionary insights specific to Madras hedgehog.

Mitochondrial DNA (mtDNA) with its rapid mutation rates and maternal inheritance, is an invaluable tool in phylogenetic studies. It has been used extensively to examine the evolutionary relationships among species within vertebrate lineages, particularly in groups like hedgehogs (Meredith et al. 2011; Bernt et al. 2013). The mitochondrial genome provides insights into species conservation, phylogeny, and evolutionary divergence (Rubinoff 2006; Ghildiyal et al. 2023). However, studies on hedgehog phylogenetics have largely focused on a limited number of species, leaving a substantial gap in our understanding of the genetic diversity within the genus. (Zeng et al. 2024) identified potential cryptic diversity within Paraechinus, emphasizing the need for further research into its genetic structure and evolutionary dynamics.

In this study, we address this gap by presenting a detailed annotation and comparative analysis of the mitochondrial genome of the Madras hedgehog. We provide a publicly available annotated mitochondrial genome for this species, offering a valuable resource for future research. Through high-throughput sequencing and comprehensive bioinformatic analysis, we provide an in-depth comparison of its mitochondrial genome, focusing on key regions such as protein-coding genes, the control region, tRNAs, and rRNAs, in relation to other species within the Erinaceidae family. By filling the genetic knowledge gap, we aim to inform future conservation efforts and provide crucial genetic resources for studying hedgehog diversity in South Asia. Additionally, we provide a detailed comparative analysis of the mitochondrial genomic data of *P. nudiventris* with closely related species, including closely related *Paraechinus* species, focusing on specific features such as base composition and gene comparisons. While previous studies have included *P. nudiventris* in phylogenetic analyses, our study emphasizes detailed annotation and comparative insights that contribute to understanding the genetic basis of evolutionary divergence within this species. By utilizing samples from opportunistic sources such as roadkill, this study is an example for the practical use of non-invasive sampling to generate valuable genetic data. The publicly available annotated mitochondrial genome for *P. nudiventris* provided here serves as a key resource for future research on hedgehog diversity and conservation in South Asia.

## Materials and Methods

### Sample Collection

Fieldwork was conducted from June 2022 to May 2023 in multiple areas across Tamil Nadu, India, specifically in Theri Kaadu, Tuticorin, Satankulam, Tirunelveli, Idaiyapatti, and Madurai (Fig. 1). A systematic random line transect survey (Hedley and Buckland, 2004) was employed across a 1×1 km^2^ grid in a study area of 62 km^2^ to detect the presence of Madras hedgehogs. Intensive fieldwork was carried out in the grids of Theri Kaadu in Thoothukudi District, with the majority of surveys conducted during the nocturnal hours (early dawn and dusk), as Madras hedgehogs are nocturnal animals, increasing the likelihood of detection. The effort included surveying several areas in the study region over a period of approximately 40 days. Field sampling primarily consisted of road-killed animals and collection of the spines from the field. For road-killed animals, the kinds of tissues collected included dried skin and other preserved tissue types. Tissue samples were obtained from dead road killed hedgehogs found along roads and were preserved using chemicals such as 70% ethanol for storage. Samples were transported to the lab and stored at -20°C for further genetic analysis.

**Figure 1.**
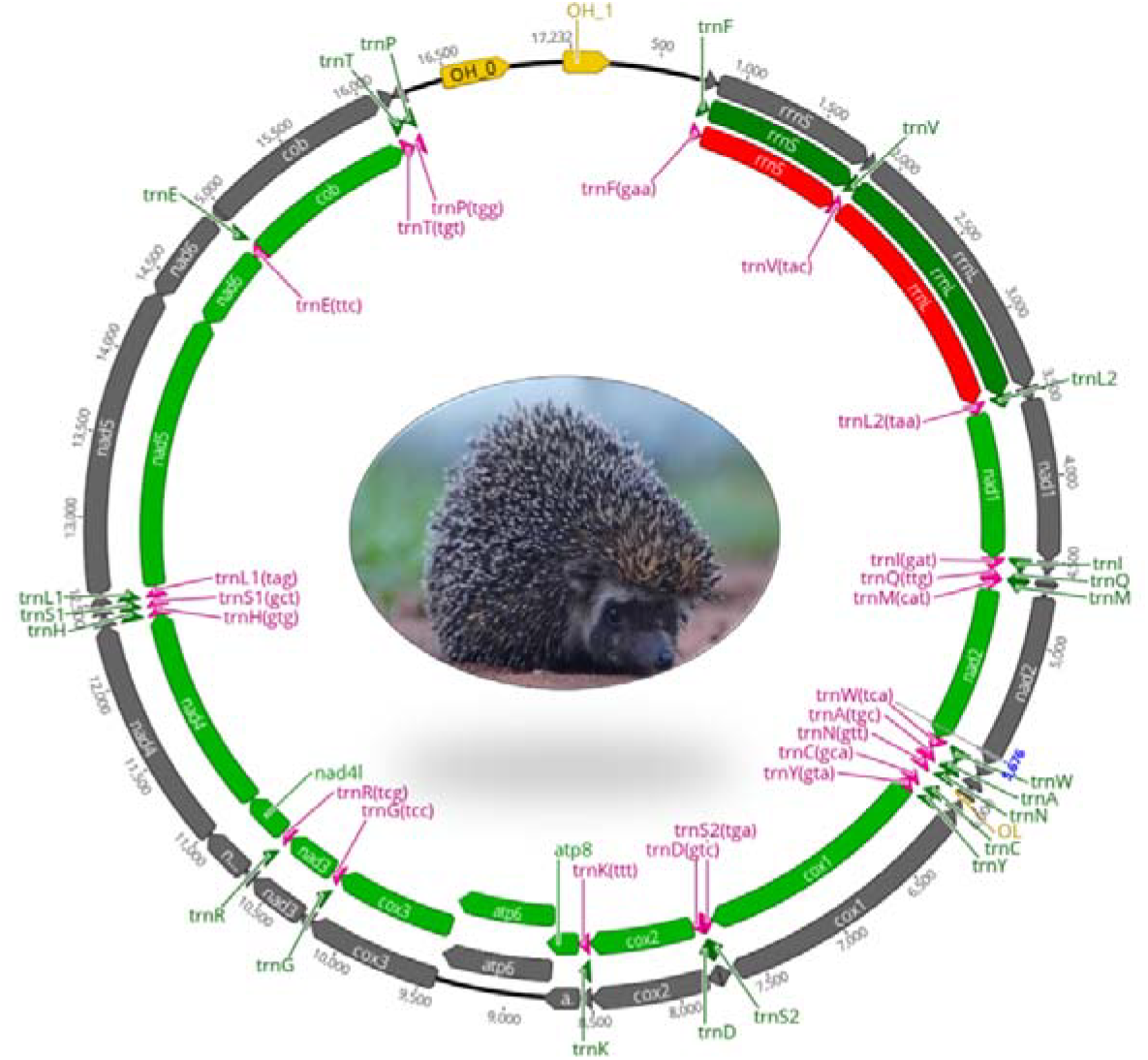
Mitochondrial genome of Madras Hedgehog (*Paraechinus nudiventris*) and Madras Hedgehog (*Paraechinus nudiventris*) from the semi-arid patches of Tiruppur District, Tamil Nadu, India.

### DNA extraction and library preparation

DNA extraction followed a standardized protocol for muscle tissues, employing SDS buffer (500 µl) as the lysis buffer and 16 µl of Proteinase K. Following overnight incubation, 3M KOAc was introduced, succeeded by a series of centrifugation steps. In two stages, the resulting supernatant was transferred to an EconoSpin column containing a binding buffer and subjected to centrifugation at 12,000 rpm. Final purification steps included an 80% ethanol wash followed by elution utilising 10 mM Tris buffer (pH 8.3). Following extraction, DNA quality was assessed using a 1.5% agarose gel. The purification method employed magnetic beads (MagSi-DNA beads-MagnaMedics) and a magnetic separator SL-MagSep96 (Steinbrenner, Germany) with an optimized MagSi-DNA Vegetal kit protocol. The library preparation followed by sequencing was carried out in an Illumina MiSeq machine at the Genomics Service Unit at Ludwig Maximillian Universitat, München, Germany.

### Mitochondrial Genome Assembly and Annotation

Sequencing data in fastQ format obtained from low-coverage Illumina whole-genome sequencing of the Madras Hedgehog (*P. nudiventris*) were used for mitochondrial genome assembly. The input data included paired-end reads with a read length of 151 base pairs (bp) and an insert size of 300 bp. Fasta files of specific mitochondrial gene sequences, including cytochrome b (cyt b), 12S rRNA, 16S rRNA, D-loop, and CO1, were selected as seeds for de novo assembly using NOVOPlasty v4.3.3 (Dierckxsens et al. 2016). The assembled mitochondrial genome was subjected to quality control and validation steps. First, the circular representation of the generated mitogenome was plotted using the CGView Server with default parameters (Grant and Stothard 2008). Additionally, the contig was confirmed using the MITOS v806 online web server (Donath et al. 2019), which provided insights into the arrangement and direction of protein-coding genes (PCGs), transfer RNAs (tRNAs), and ribosomal RNAs (rRNAs).

### Comparative and Phylogenetic Analysis

To investigate the phylogenetic relationships among the studied taxa and estimate divergence times, a comprehensive dataset comprising 19 mitogenomes, including 18 downloaded sequences along with the newly assembled Madras Hedgehog (*P. nudiventris*), was analyzed. The mitogenome sequences of these hedgehog species were aligned with sequences from other taxa, including outgroup species for comparative purposes. The hedgehog species included in our analysis were *Atelerix albiventris* (OP654705.1) (He and Zeng, unpublished), *Mesechinus dauuricus* (OR264473.1) (Bayarlkhagva et al., unpublished), *Erinaceus europaeus* (NC_002080.2) (Krettek et al. 1995), *Erinaceus amurensis* (KX964606) (Kim et al. 2017), *Erinaceus concolor* (OP654723.1) (He and Zeng, unpublished), *Hemiechinus auritus* (AB099481.1) (Nikaido et al. 2003), *Mesechinus hughi miodon* (KT824773.1) (Kong et al., unpublished), *Paraechinus hypomelas* (OP654725.1) (He and Zeng, unpublished), *Paraechinus micropus* (OP654708.1) (He and Zeng, unpublished), *Hylomys suillus* (PRJNA927338) (Arnason et al. 2008) and *Echinosorex gymnura* (AF348079) (Lin, direct submission). Additionally, we included outgroup species *Talpa europaea* (Y19192) (Mouchaty et al. 2000), *Sorex araneus* (KT210896) (Zhu et al. 2017). These species were selected to represent a diverse range of taxa within the Erinaceidae family, with the outgroup species providing comparative context for the analysis.

The best-fit nucleotide substitution model (GTR + I + G) was determined through PartitionFinder2 (Lanfear et al., 2017) based on the lowest Bayesian information criterion (BIC). Subsequently, a maximum-likelihood (ML) tree was constructed using IQ-TREE (Minh et al. 2020) with 1000 bootstrap replicates to assess branch support. For Bayesian inference, MrBayes (Ronquist et al. 2012) was employed with settings specifying the nucleotide substitution model (NST = 6) and gamma-distributed rate variation across sites. MCMC analyses were run for 1,000,000 generations, sampling every 100 generations, with convergence assessed by standard diagnostics. Additionally, divergence times among Erinaceidae species were estimated using a Bayesian relaxed clock method in BEAST v2.4.7 (Bouckaert et al. 2019), incorporating fossil calibration points for specific nodes. Two primary calibration points were applied: (1) representing the divergence of Soricidae and Talpidae, which has been reliably dated in previous studies (Springer et al. 2003), and (2) at the divergence between the gymnures and all hedgehogs based on fossil evidence from early erinaceomorphs (Bannikova et al. 2014). A third, secondary calibration point was placed at the split between *Erinaceus concolor* and *Erinaceus amurensis*, reflecting more recent divergence events within the Erinaceus (Seddon et al. 2001; He et al. 2012). Markov chain Monte Carlo (MCMC) runs were performed for 1,000,000 generations, with effective sample size (ESS) values monitored to ensure adequate mixing and convergence. The resulting phylogenetic tree and divergence time estimates provide valuable insights into the evolutionary history of the studied taxa.

## Results

### Mitochondrial Genome Overview

Whole genome sequencing produced a total of 8,139,858 reads, of which 16,040 mitochondrial reads were aligned and 10,374 were assembled, resulting in an average organelle coverage of 143x for the Madras Hedgehog’s mitochondrial genome. The final contig length was 17,232 bp, representing the complete mitochondrial genome (Fig. 1), with a nucleotide composition of A: 34.3%, T: 34.2%, G: 12.2%, and C: 19.3%, resulting in a GC content of 31.5%. The genome contains the standard set of 13 protein-coding genes (*nad1, nad2, cox1, cox2, atp8, atp6, cox3, nad3, nad4l, nad4, nad5, nad6, cob*), 22 transfer RNAs (tRNAs), and 2 ribosomal RNA (rRNA) genes (Table 1).

**Table 1.**
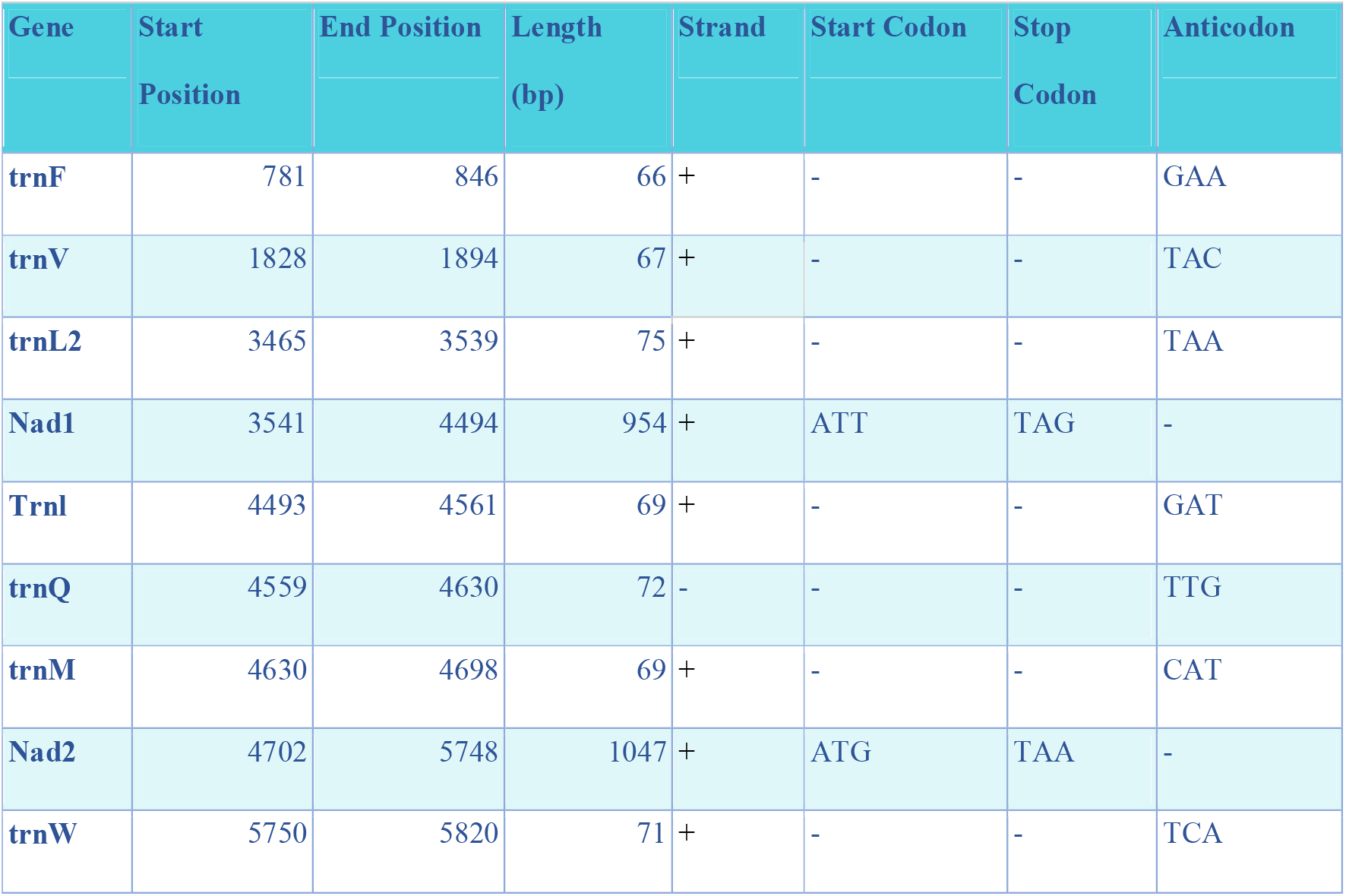
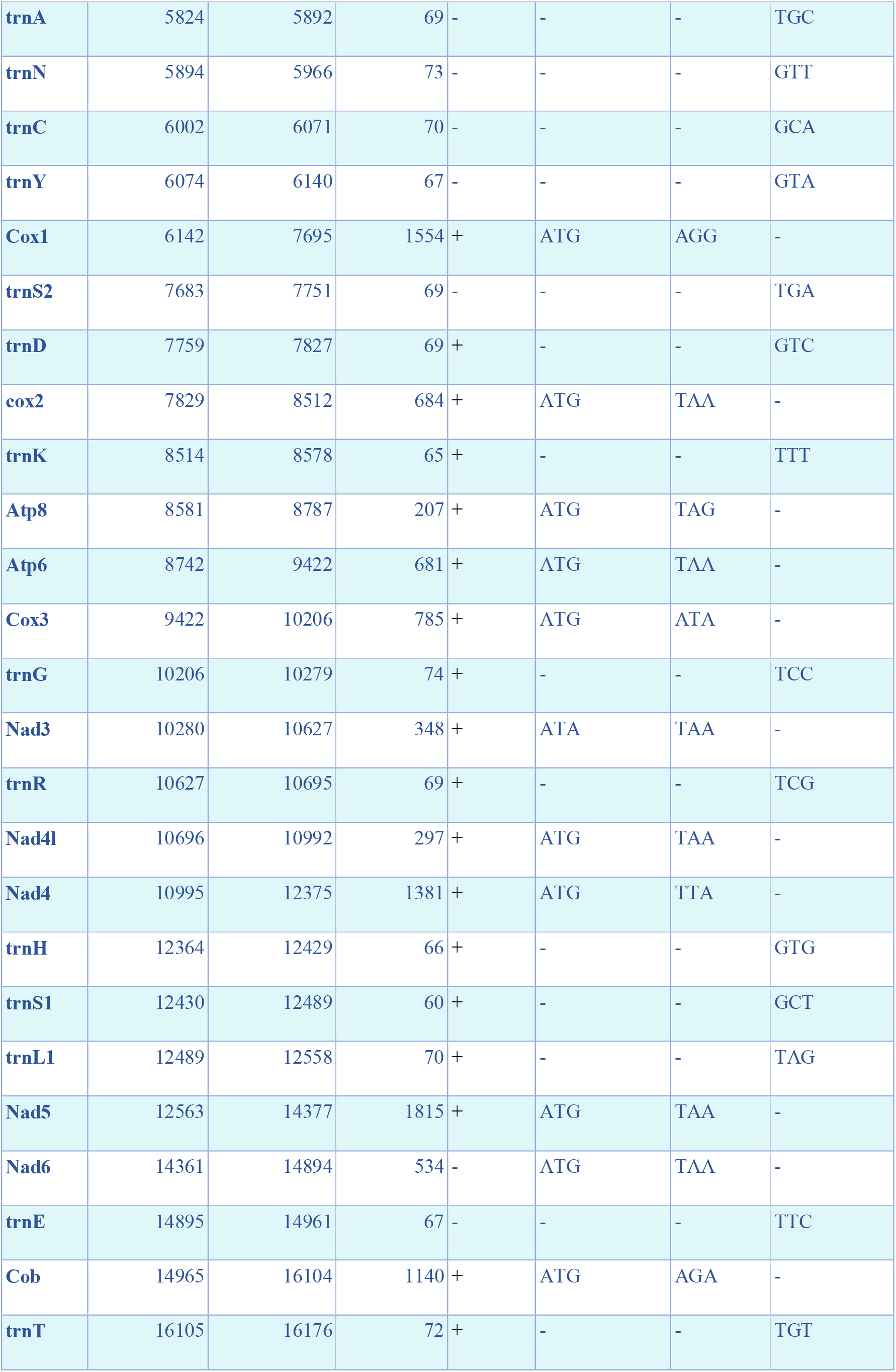
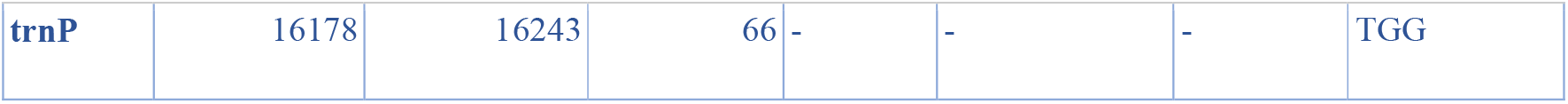
Details of the annotated genes.

### Base Composition and Nucleotide Skewness

Across the five hedgehog species analyzed, the mitochondrial genomes demonstrate a distinct A+T bias, which is a characteristic feature of vertebrate mitochondrial genomes. The combined adenine (A) and thymine (T) content ranged from 63.8% in *Erinaceus europaeus* to 69.1% in *Hemiechinus auritus*. The AT-skew values for all species were close to zero, indicating a relatively balanced ratio of adenine to thymine. *Erinaceus europaeus* exhibited a slightly negative AT-skew (−0.12), while GC-skew values were consistently negative, ranging from - 0.14 in *Erinaceus europaeus* to -0.25 in *Erinaceus amurensis*. The non-coding regions of the Madras Hedgehog’s mitochondrial genome included typical tRNA clusters and the origins of heavy strand replication (*OH_0* and *OH_1*), located between *tRNA-P* and *tRNA-F*. The secondary structure of all tRNAs followed the classic cloverleaf model, except for *tRNA-Ser*, which exhibited an atypical structure. The control region (D-loop), a 1,732 bp non-coding segment situated between *tRNA-Phe* and *tRNA-Pro*, was identified based on its location and sequence similarity to other known D-loops in vertebrate mitochondrial genomes.

### Protein-coding Genes (PCGs)

The total length of the protein-coding genes (PCGs) across the hedgehog species ranged from 11,404 bp in *Erinaceus amurensis* to 11,427 bp in *P. nudiventris*. The base composition of the PCGs showed an A+T bias, with A+T content ranging from 67.6% in *Erinaceus europaeus* to 69.5% in *P. nudiventris*. The AT-skew values for the PCGs were negative ranging from -0.046 in *Erinaceus amurensis* to -0.068 in *P. nudiventris*. Similarly, the GC-skew values were negative, ranging from -0.22 in *P. nudiventris* to -0.28 in *Erinaceus amurensis*.

### Ka/Ks Ratios for Protein-coding Genes

Ka/Ks ratios were calculated for the PCGs using MEGA, and the values ranged from 0.033 to 0.39. The highest Ka/Ks ratio was observed for *atp8* (0.39) while the lowest ratios were observed in the cytochrome oxidase genes (*cox1, cox2*, and *cox3*), with values ranging from 0.033 to 0.1. Intermediate Ka/Ks values observed in the NADH dehydrogenase subunits (*nad2, nad5*).

### Ribosomal RNA (rRNA) Genes

The total length of the rRNA genes was consistent across the species, ranging from 2,513 bp in *Erinaceus amurensis* to 2,560 bp in *Mesechinus hughi*. The A+T content ranged from 66.2% in *Mesechinus hughi* to 66.9% in *Hemiechinus auritus*. The AT-skew values were positive, ranging from 0.146 in *Mesechinus hughi* to 0.154 in both *P. nudiventris* and *Hemiechinus auritus*, while GC-skew values were slightly negative, ranging from -0.025 to -0.045.

### Control Region

The control region (CR) varied in length from 1,362 bp in *Mesechinus hughi* to 1,988 bp in *Erinaceus europaeus*. The A+T content ranged from 65.9% in *Erinaceus europaeus* to 72.9% in *Hemiechinus auritus*. Using Tandem Repeats Finder (Benson, 1999), we identified two tandem repeat regions in *P. nudiventris* between positions 943–1,193 and 1,190–1,222. The first repeat (250 bp) contained the motif “ACGCAT” with a copy number of 41.8, while the second (32 bp) consisted of the motif “CGCATA” with a copy number of 5.5.

**Table 2.** Comparative analysis of Madras hedgehog with European, Amur, Hugh and long-eared hedgehogs for different aspects of the mitochondrial genome.

| Feature/Species | Size (bp) | A% | T% | G% | C% | G+C% | AT-skew | GC-skew |
| --- | --- | --- | --- | --- | --- | --- | --- | --- |
| **Complete genome** |  |  |  |  |  |  |  |  |
| *Paraechinus nudiventris* | 17232 | 34.3 | 34.2 | 12.2 | 19.3 | 31.5 | 0.002 | -0.22 |
| *Erinaceus europaeus* | 17447 | 28 | 35.8 | 15.5 | 20.7 | 36.2 | -0.12 | -0.14 |
| *Hemiechinus auritus* | 17283 | 35.2 | 33.9 | 11.7 | 19.2 | 30.9 | 0.019 | -0.24 |
| *Mesechinus hughi* | 16842 | 34.9 | 33.5 | 11.9 | 19.7 | 31.6 | 0.019 | -0.24 |
| *Erinaceus amurensis* | 16941 | 33.9 | 32.6 | 12.4 | 21.1 | 33.5 | 0.019 | -0.25 |
| **Protein-coding genes** |  |  |  |  |  |  |  |  |
| *Paraechinus nudiventris* | 11427 | 32.4 | 37.1 | 11.8 | 18.7 | 30.5 | -0.068 | -0.22 |
| *Erinaceus europaeus* | 11405 | 31.5 | 36.1 | 12.1 | 20.2 | 32.4 | -0.067 | -0.25 |
| *Hemiechinus auritus* | 11408 | 33 | 36.5 | 11.4 | 19.1 | 30.5 | -0.049 | -0.25 |
| *Mesechinus hughi* | 11412 | 32.9 | 36.3 | 11.5 | 19.4 | 30.9 | -0.049 | -0.25 |
| *Erinaceus amurensis* | 11404 | 31.8 | 34.9 | 11.9 | 21.4 | 33.3 | -0.046 | -0.28 |
| **tRNAs** |  |  |  |  |  |  |  |  |
| *Paraechinus nudiventris* | 1448 | 34.3 | 33.2 | 17.7 | 14.7 | 32.5 | 0.016 | 0.091 |
| *Erinaceus europaeus* | 1517 | 34.7 | 32.1 | 17.7 | 15.5 | 33.2 | 0.038 | 0.067 |
| *Hemiechinus auritus* | 1514 | 34.5 | 32.5 | 18 | 14.9 | 33 | 0.03 | 0.094 |
| *Mesechinus hughi* | 1413 | 34.6 | 31.9 | 18.2 | 15.3 | 33.5 | 0.041 | 0.088 |
| *Erinaceus amurensis* | 1515 | 34.1 | 32 | 17.9 | 16 | 33.9 | 0.031 | 0.056 |
| **rRNAs** |  |  |  |  |  |  |  |  |
| *Paraechinus nudiventris* | 2543 | 37.8 | 27.6 | 16.9 | 17.7 | 34.6 | 0.154 | -0.025 |
| *Erinaceus europaeus* | 2534 | 37.1 | 27.3 | 17 | 18.5 | 35.6 | 0.152 | -0.04 |
| *Hemiechinus auritus* | 2558 | 38.2 | 28 | 16.1 | 17.7 | 33.8 | 0.154 | -0.04 |
| *Mesechinus hughi* | 2560 | 37.7 | 28.1 | 16.3 | 17.9 | 34.2 | 0.146 | -0.045 |
| *Erinaceus amurensis* | 2513 | 37.3 | 27.5 | 16.8 | 18.3 | 35.2 | 0.151 | -0.044 |
| **Control Region** |  |  |  |  |  |  |  |  |
| *Paraechinus nudiventris* | 1769 | 35 | 32.4 | 11.7 | 20.8 | 32.5 | -0.038 | -0.28 |
| *Erinaceus europaeus* | 1988 | 37.8 | 32.9 | 12.1 | 17.2 | 29.3 | 0.069 | -0.175 |
| *Hemiechinus auritus* | 1813 | 38.6 | 34.3 | 9.3 | 17.8 | 27.1 | 0.059 | -0.311 |
| *Mesechinus hughi* | 1362 | 36.7 | 33 | 10.8 | 19.5 | 30.3 | 0.053 | -0.288 |
| *Erinaceus amurensis* | 1509 | 34.7 | 33.1 | 12.1 | 19.3 | 31.5 | 0.023 | -0.23 |

### Phylogenetic analysis

Bayesian inference and maximum likelihood analyses were used to construct phylogenetic trees, both of which produced identical topologies. Strong statistical support was observed for intergenic relationships within Erinaceinae, highlighting the distinct evolutionary lineages (Figure 2a). Divergence time estimates, based on three calibration points, suggest that the most recent common ancestor (MRCA) of Galericinae existed approximately 39.6 mya, while Erinaceinae emerged around 30.1 mya (Figure 2b). The divergence of the Erinaceinae lineage during the Miocene in the Neogene period (approximately 15.9 mya) underscores its evolutionary significance. Notably, the Madras hedgehog split from *P. micropus*, another Indian hedgehog species, during the Pleistocene epoch approximately 3.69 mya.

**Figure 2.**
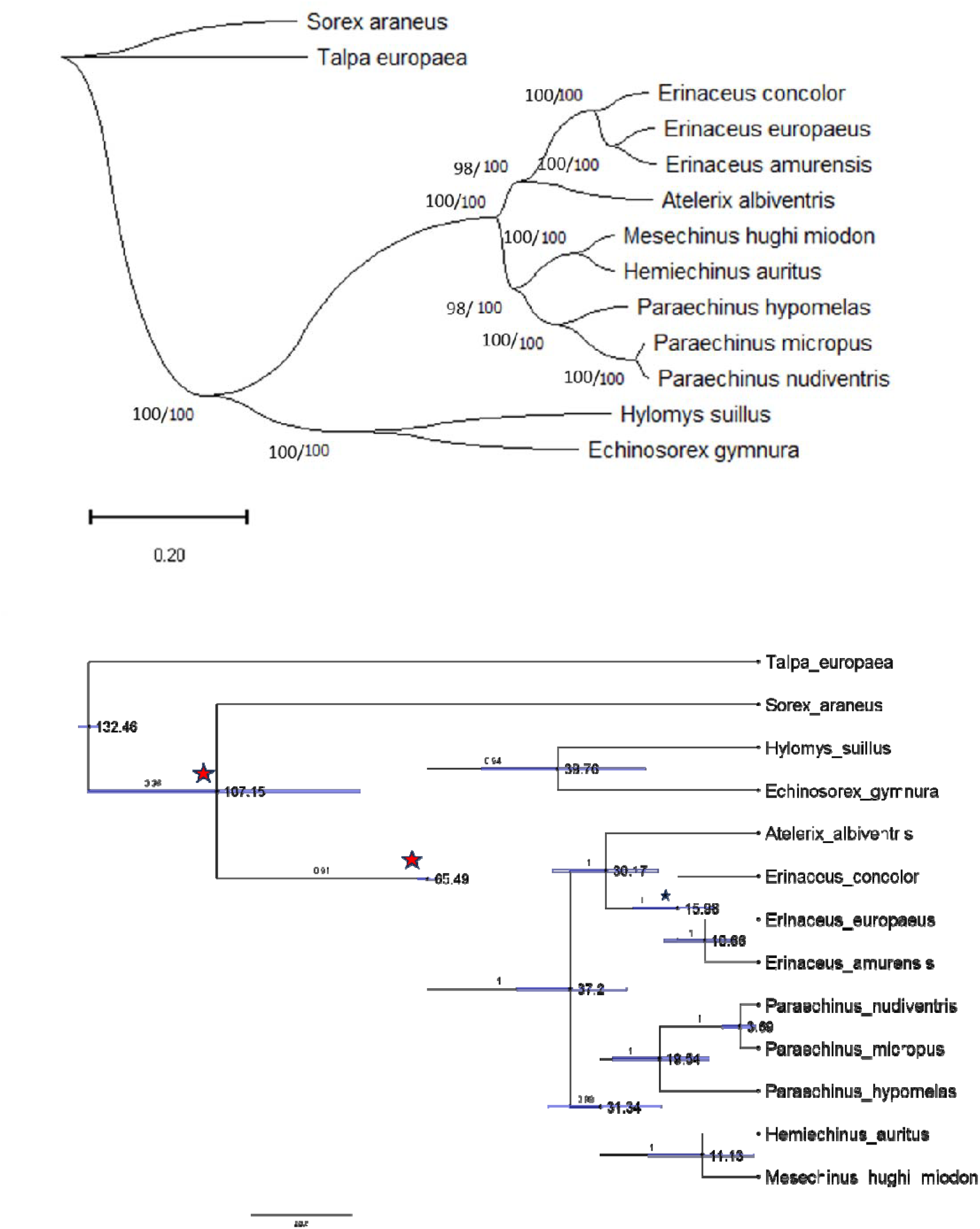
a) Bayesian tree of molecular phylogeny of the Madras hedgehog with other species in the Erinaceinae and Galericinae families. The bootstrap values of the maximum-likelihood trees are shown before the values from the Bayesian analyses separated by a ‘/’. b) Calibrated phylogenetic tree generated using BEAST v1.10.4 using two primary calibration points (red stars) and a secondary point (marked by black star). The blue bars indicate the 95% highest probability density (HPD) intervals surrounding the mean divergence estimates.

## Discussion

The mitochondrial genome of the Madras Hedgehog presents an opportunity to expand our understanding of the evolutionary dynamics within the Erinaceidae family. The distinct A + T bias observed across hedgehog species aligns with well-documented trends in vertebrate mitochondrial genomes, reflecting mutational pressures and replication dynamics. Negative GC-skew values across all species, particularly in *Erinaceus amurensis*, further emphasize species-specific compositional biases that may influence gene expression and replication efficiency. Phylogenetic analyses revealed robust relationships within Erinaceinar, with both Bayesian and maximum likelihood methods yielding consistent topologies. The Madras Hedgehog emerged as a distinct lineage, diverging from *P. micropus* during the Pleistocene epoch. This timing aligns with significant climatic and geological changes that likely influenced speciation and dispersal patterns (Cailleux et al. 2023). The estimated divergence of Erinaceinae during the Miocene suggests an earlier phase of diversification within this subfamily, likely shaped by the changing landscapes of the Neogene period (Zhao et al. 2012). Divergence time estimates provide a framework for understanding the evolutionary trajectory of Erinaceinae. The MRCA of Galericinae, estimated at 39.6 mya, underscores the deep evolutionary roots of this family, while the emergence of Erinaceinae approximately 30.1 mya highlights its subsequent diversification. The split between the *P. nudiventris* and *P. micropus* during the Pleistocene points to more recent ecological and geographical influences driving speciation.

The protein-coding genes exhibit conserved lengths and A+T-rich composition, indicative of functional constraints. Elevated Ka/Ks rations for atp8 highlight its evolutionary flexibility, potentially reflecting adaptive responses, while the strong purifying selection in cytochrome oxidase genes underscores their critical role in oxidative phosphorylation. These findings align with broader patterns of mitochondrial gene evolution in vertebrates.

Variations in the control region, including tandem repeats in *P. nudiventris*, suggest functional adpatations in mitochondrial replication and transcription regulation. The identified repeat motifs and conserved sequence blocks emphasize the regulatory significance of this region. Ribosomal RNA gene stability further supports their role in maintaining essential mitochondrial functions, with slight variations hinting at lineage-specific evolutionary pressures. This study integrates findings on mitochondrial genome composition, gene evolution, and regulatory regions, providing a comprehensive view of the Madras hedgehog’ mitochondrial genome. These insights contribute to a broader understanding of genetic diversity and evolutionary trajectories within the Erinaceidae family.

## Acknowledgements

All institutional and/or national guidelines and regulations regarding the collection of specimens were followed. Special thanks to National Biodiversity Authority (NBA), Government of India for their approval and permission for transfer of Madras Hedgehog samples for research as per the Biological Diversity Act, 2002 and Rule 14 of the Biological Diversity Rules. We would like to acknowledge the support from NBA for transferring the samples to Vienna. (File number: NBA/Tech Appl/9/lNBA1202203611/22/22-23/3041). Further, thanks to Tamil Nadu Biodiversity Board (TNBB) (Ref No.TNBB/1357/2022/B3,14.02.2022) and Tamil Nadu state forest department Chennai for granting us to conduct this research study (Proceeding No.WL5 (A)31710/2021-Permission No: 92/2022). Thanks to the Internationally Recognized Certificate of Compliance - (ABSCH-IRCC-IN-264698-1) from the ABS house. This work was carried out when BK worked with Indian Institute of Science Education and Research (IISER), Tirupati, India under the NPDF fellowship. Thanks to Nandini Rajamani from IISER Tirupati for her encouragement to carry out this work. This work was made possible thanks to a DST – SERB National Post-Doctoral Fellowship (PDF/2019/003178) and IISER Tirupati institutional support. BK field work is supported by the TAAL Tech India Private Limited, Pune, India.

## Author contributions

B.K. and H.M. designed the study; B.K. supervised the study in collaboration with H.M; B.K carried out the field work and B.K and T.V handled permits for samples exchange. T.V. processed the Madras Hedgehog sequencing and assembly; T.V. compiled and analyzed the data; B.K. and T.V. drafted the manuscript; All authors revised the manuscript.

## CRediT authorship contribution statement

**Thapasya Vijayan:** Data curation, Investigation, Methodology, Formal Analysis, Validation, Visualization, Writing – Original draft, review & editing.

**Harald Meimberg:** Resources, Investigation, Methodology, Conceptualization, Supervision, Writing – review & editing, Funding acquisition.

**Brawin Kumar**: Supervision, Conceptualization, Investigation, Data Curation, Validation, Fund Raising, Writing – original draft, Writing – review & editing.

## Data availability

The assembles genome is available at NCBI.

